# Deciphering the targets of retroviral protease inhibitors in *Plasmodium berghei*

**DOI:** 10.1101/334524

**Authors:** Noah Machuki, Reagan Mogire, Loise Ndung’u, Peter Mwitari, Francis Kimani, Damaris Matoke-Muhia, Daniel Kiboi, Gabriel Magoma

## Abstract

Retroviral protease inhibitors (RPIs) such as lopinavir (LP) and saquinavir (SQ) are active against *Plasmodium* parasites. However, the exact molecular target(s) for these RPIs in the *Plasmodium* parasites remains poorly understood. We hypothesised that LP and SQ suppress parasite growth through inhibition of aspartyl proteases. Using reverse genetics approach, we embarked on separately generating knockout (KO) parasite lines lacking Plasmepsin 4 (PM4), PM7, PM8, or DNA damage-inducible protein 1 (Ddi1) in the rodent malaria parasite *Plasmodium berghei* ANKA. We then tested the suppressive profiles of the LP/Ritonavir (LP/RT) and SQ/RT as well as antimalarials; Amodiaquine (AQ) and Piperaquine (PQ) against the KO parasites in the standard 4-day suppressive test. The Ddi1 gene proved refractory to deletion suggesting that the gene is essential for the growth of the asexual blood stage parasites. Our results revealed that deletion of PM4 significantly reduces normal parasite growth rate phenotype (*P* = 0.003). Unlike PM4_KO parasites which were less susceptible to LP and SQ (*P* = 0.036, *P* = 0.030), the suppressive profiles for PM7_KO and PM8_KO parasites were comparable to those for the WT parasites. This finding suggests a potential role of PM4 in the LP and SQ action. On further analysis, modelling and molecular docking studies revealed that both LP and SQ displayed high binding affinities (-6.3 kcal/mol to -10.3 kcal/mol) towards the *Plasmodium* aspartyl proteases. We concluded that PM4 plays a vital role in assuring asexual stage parasite fitness and might be mediating LP and SQ action. The essential nature of the Ddi1 gene warrants further studies to evaluate its role in the parasite asexual blood stage growth as well as a possible target for the RPIs.

**Author summary:** The antiretroviral drugs (ARVs) such as LP or SQ that inhibit viral proteases reduce the rate of multiplication of the malaria parasites. The mode of action of these drugs against the parasites is however poorly understood. The proteases are among the enzymes that play essential roles in *Plasmodium* parasites. We sought to investigate the possible mode of action of these drugs by generating mutant parasites lacking specific aspartyl proteases namely PM4, PM7, PM8 or Ddi1 and then evaluate the susceptibility of the mutants to LP and SQ. We successfully generated parasites lacking either PM4, PM7 or PM8 but Ddi1 gene was refractory to deletion. From our data, we demonstrate that, unlike PM7 and PM8, the PM4 and Ddi1 are essential enzymes for asexual blood stage parasite fitness and survival and that the PM4 might be a target for the viral protease inhibitors in reducing parasite growth and multiplication. Further experiments using molecular docking tools show that LP or SQ have a high binding affinity for the *Plasmodium* aspartyl proteases.

## Introduction

Notwithstanding the immense investments in malaria control programs to date, it remains to be a significant global health problem in most regions of the world including Africa, Asia and parts of the Eastern Mediterranean Region (1,2). The sub-Saharan part of Africa continues to bear the highest burden of the disease with over 90% of the cases occurring in this region, especially in children under five years of age. In the year 2016 alone, an estimated 285 000 children succumbed to malaria in Africa (2).

The emergence and spread of resistance to available drugs including the artemisinin-based combination therapies (ACTs) have aggravated the burden of the malaria disease. Incidences of parasite resistance to the ACTs were first reported in western Cambodia and currently slowly spreading to other parts of Asia. The South East Asia region occupies a historical record as a site of emerging resistance to the previous first-line antimalarial therapies which later rapidly spread across the African countries where malaria transmission is consistently high (3–6). Since the options of drugs for which the human malaria parasite *Plasmodium falciparum* has not evolved resistance is rapidly diminishing, new and rational approaches to the prevention and treatment of malaria infections are urgently needed.

The burden of malaria is compounded with HIV/AIDS infections which are also concentrated in the malaria-endemic regions, primarily sub-Saharan Africa. This geographical overlap has raised opportunities and concerns for potential immunological, social, therapeutic and clinical interactions (7). Previous studies have demonstrated that the antiretroviral therapy, especially RPIs exert a potent effect against both the drug-sensitive and drug-resistant *P. falciparum* (8–14), as well as a reduction in the incidence of malaria (15). For instance, seven RPIs inhibit the development of *P. falciparum* parasites in vitro with lopinavir yielding moderate synergy with lumefantrine (12). The RPIs are typical examples of drugs that target an aspartyl protease in HIV, HIV-1 aspartyl protease (16,17). Like in HIV, aspartyl proteases play essential roles in the biology of *Plasmodium* parasites and thus are “druggable” targets (18–21). The blood stage aspartyl proteases are responsible for host haemoglobin digestion, rupturing of the host erythrocytes (22), as well as processing of effector proteins for export to the infected erythrocytes (23). The RPIs are predicted to exert their antimalarial activity in the blood stages of parasite life cycle (8,13). Previous studies geared towards understanding the mode of action of the RPIs in suppressing the growth of *Plasmodium* parasites focused on pepsin-like proteases (PMs) even though *Plasmodium* species express a retropepsin-like protease, referred to as Ddi1 (24).

Using the rodent malaria parasite, *P. berghei*, and highly efficient *Plasmo*GEM genetic modification vectors, we engineered the *Plasmodium* aspartyl proteases; PM4, PM7, PM8 and Ddi1 in our quest to understand the possible mechanisms of action of LP and SQ (the most active RPIs). Here, we report that the PM4 and Ddi1 genes are essential for asexual blood stage parasite, but PM7 and PM8 genes are not. We further discuss the growth rate phenotypes of the KO parasites lacking PM7, PM8 or PM4 genes as well as the *in vivo* susceptibility profiles of the KO parasites to LP and SQ. Finally, using modeling and molecular docking, we predict the *in silico* binding affinities of the LP and SQ towards PM4, PM7, PM8 or Ddi1. The findings reveal that PM4 assures parasite fitness in the asexual stage, mediates the possible mechanism of action of LP and SQ in parasite growth suppression as well as the refractory nature of Ddi1. Here, we for the first time provide evidence that the retropepsin-like protease is essential for the asexual stage of the malaria parasite.

## Materials and Methods

### Parasites, vectors, hosts and test compounds

A transgenic *P. berghei* ANKA strain 676m1cl1{PbGFP-LUC(con)} which expresses a fusion protein GFP-Luciferase (25), maintained in cryopreserved stocks in Kenya Medical Research Institute (KEMRI), was used in this study. The KO vectors (PbGEM-039254, PbGEM- 086320, PbGEM-057101, and PbGEM-097527) were obtained under a material transfer agreement with the PlasmoGEM project at the Wellcome Trust Sanger Institute (PG-MTA-0093). Naive male Swiss albino mice (6-7 weeks old), weighing 18-20g were acquired from the KEMRI animal house and used as models in this study. The animals were kept in the KEMRI animal house in standard polypropylene cages and fed on commercial rodent feed and water *ad libitum*. LP, SQ and RT were purchased from Sigma Aldrich (USA). All the test compounds were prepared by solubilising them in a solution consisting of 70% Tween-80 (density=1.08gml^-1^) and 30% ethanol (density=0.81 gml^−1^) and were diluted ten times with distilled water. The diluted Tween-80 and ethanol solution was used as a vehicle and control for the drug profile assays.

### Ethics Statement

All protocols were conducted in accordance with prior approvals obtained from the Kenya Medical Research Institute (KEMRI)’s Scientific Ethical Review Unit (SERU; 3572) and the KEMRI Animal Care and Use Committee (ACUC). The KEMRI ACUC adheres to national regulations on the care and use of animals in research in Kenya enforced by the National Commission for Science, Technology and Innovation (NACOSTI). The Institute has a Public Health Service (PHS)-approved assurance, number F16-00211 (A5879-01) from the Office of Laboratory Animal Welfare (OLAW) and commits to the International Guidelines for Biomedical Research Involving Animals.

### Generation of the knockout parasite lines

#### Isolation, digestion, and purification of vector DNA

Plasmid DNAs were isolated from overnight cultures of *E. coli* (vector hosts) in terrific broth (TB) supplemented with kanamycin using QIAfilter Plasmid Midi Kit. The isolated plasmid was prepared for transfection by Not1 restriction digestion to liberate the vector backbone and was purified using standard ethanol precipitation. Each isolated construct was dried at 65°C for 5 min, dissolved in 10 µl of PCR water and the plasmid DNA concentration determined in a 1μl sample volume using a NanoDrop. Both restriction digestion and diagnostic polymerase chain reaction (PCR), using QCR2/GW2 primer pair, were used as vector verification strategies.

#### Collection of the *P. berghei* schizonts for transfection

Collection of the *P. berghei* schizonts for transfection was done using standard protocols as described by (26). Briefly, for each of the vectors, at least three mice were used for schizont culture. *P. berghei* parasites were propagated by intraperitoneal (IP) injection into the mice and were harvested for schizont culture at 3% parasitaemia using cardiac puncture. The asexual stage parasites (rings) were cultured in vitro at 37°C in tightly sealed and gassed flasks containing 100ml of schizont culture medium (72 ml RPMI1640, 25 ml freshly thawed FBS, 1ml of antibiotic Pen/Strep (1:100 Penicillin/streptomycin) and 2ml 0.5M NaHC03). Each flask contained 2ml of blood and was incubated overnight, in a shaker incubator. The schizonts were harvested after 22 hours, purified by Nycodenz density gradient centrifugation and each pellet re-suspended in schizont culture media, ready for transfection.

#### Transfection, selection and genotype analysis of mutant parasites

For every transfection, 20μl of schizont pellet was used. Parasite pellet was re-suspended in 100μl AMAXA supplemented nucleofector solution containing purified vector DNA. Exactly 100μl of the solution (vector DNA /nucleofector solution and schizonts) were pipetted into the Amaxa cuvette. The electroporation was done using program U33 of the Nucleofector 2B Device (Lonza). The electroporated parasites were injected intravenously (IV) into mice (two mice per transfection). Twenty-four hours post parasite inoculation, pyrimethamine drug (7 mg/mL), in drinking water, was provided to the mice for nine days. To generate a genetically homogenous parasite, the KO parasites were diluted to approximately 0.1%, passaged successively three times in naïve mice each time under the Pyrimethamine selection pressure. After cloning the recovered KO parasites, the genomic DNA (gDNA) was extracted and purified using QIAamp DNA Mini Kit following instructions from the manufacturer. The diagnostic and genotyping PCR analysis was used to genotype the KO lines using three primer pairs; QCR2/GW2, GT/GW1 or GW2 and QCR1/QCR2. QCR1 and QCR2 anneal to the genomic sequence of the targeted locus, GT anneals to the genomic sequence outside of but preceding the region covered by the gDNA clone, while GW1/2 anneals to the sequence of the selectable marker cassette (human dihydrofolate reductase; hDHFR), (Table 1A, 1B and 1C). The PCR amplification was done using 2xGoTag Green master mix for thirty cycles with an annealing temperature of 50°C and an extension temperature of 62°C. The PCR amplicons were analysed on a 1% agarose gel electrophoresis.

**Table (A):**
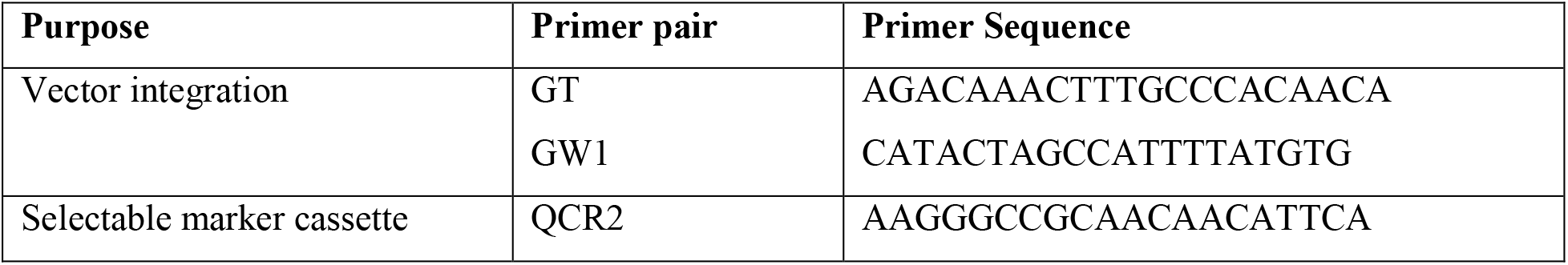
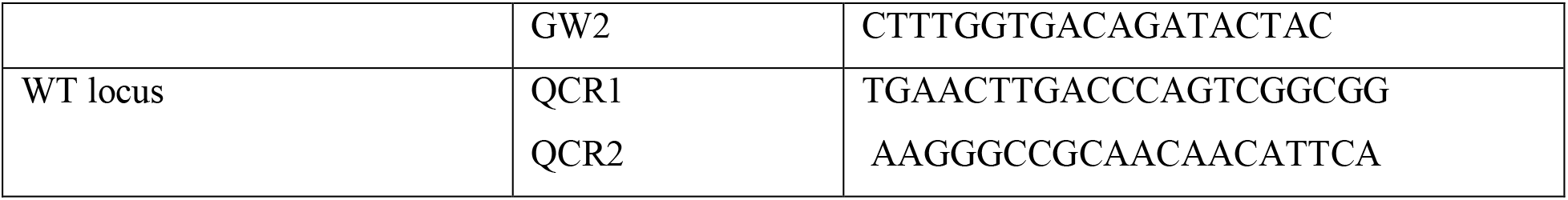
Primer pairs used in PCR amplification of gDNA isolated from PM4_KO parasites

**Table (B):**
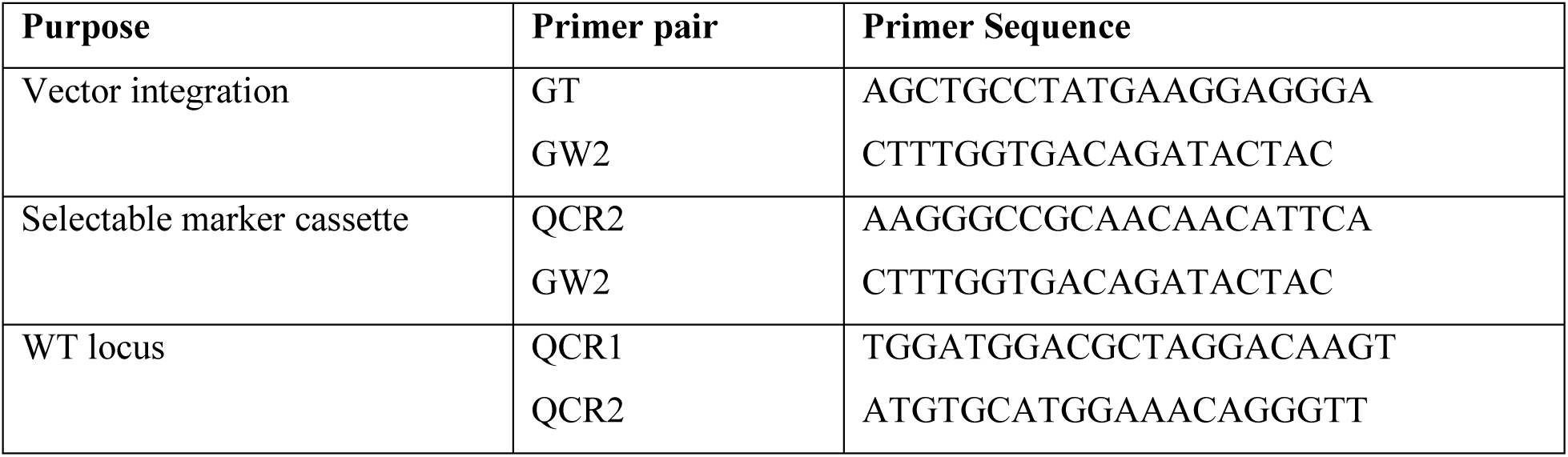
Primer pairs used in PCR amplification of gDNA isolated from PM7_KO parasites

**Table (C):**
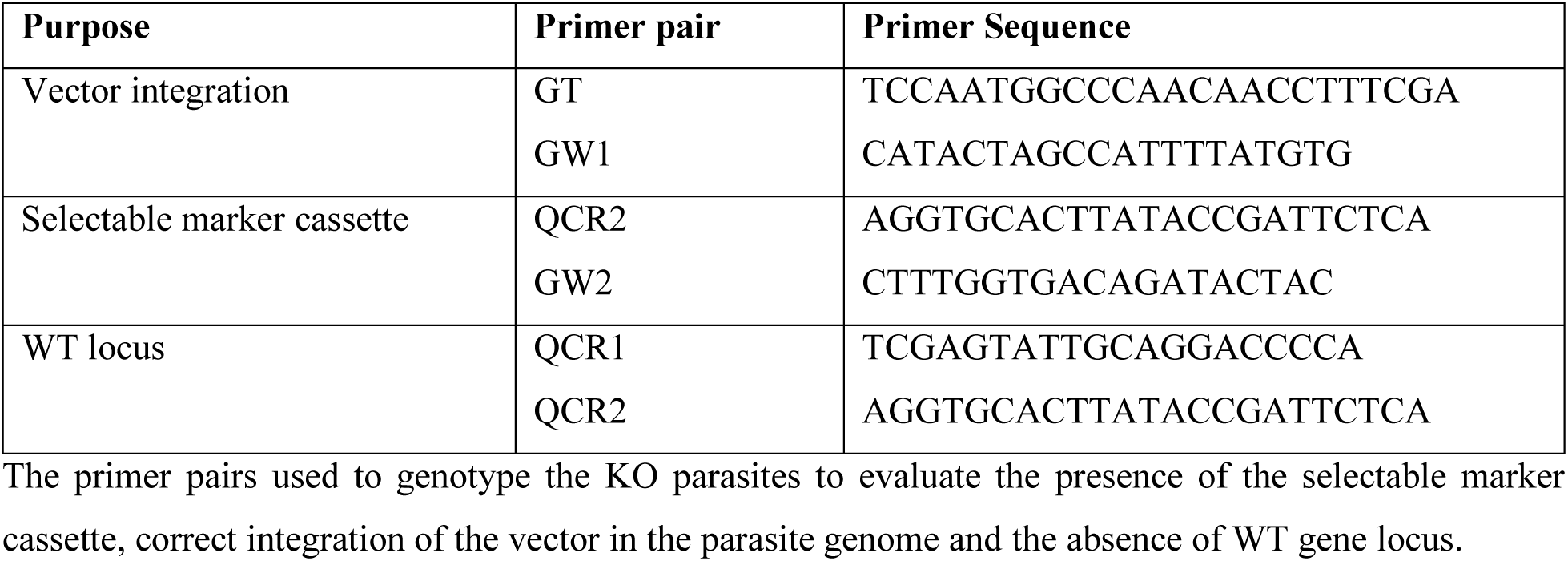
Primer pairs used in PCR amplification of gDNA isolated from PM8_KO parasites

The primer pairs used to genotype the KO parasites to evaluate the presence of the selectable marker cassette, correct integration of the vector in the parasite genome and the absence of WT gene locus.

#### Drug sensitivity profiles

Drug sensitivity profiles were determined using the quantitative standard 4-Day suppressive test (27), after four consecutive drug dosages. Briefly, for LP/RT and SQ/RT, groups of eight mice were used (four mice for each mutant parasite line and the other four for the WT parasite). For AQ or PQ, groups of sixteen mice were used (eight mice for mutant parasite line and the other eight for the WT parasites for two different AQ and PQ dosages). Each control group had four mice. Each mouse was IP inoculated with 200µl of 1× 10^5^ parasitised erythrocytes from infected donor mice, on the first day (day 0; D0). Two hours after parasite inoculation, four mice from the groups received 0.2ml of LP/RT; 40mg/kg.10mg/kg, SQ/RT; 50mg/kg.5mg/kg, AQ; 2.5 mg/kg, AQ; 1.25mg/kg, PQ; 2.5mg/kg and PQ; 1.25mg/kg, once daily oral gavage. The drug treatment proceeded for three consecutive days. On the fifth day (day 4: D4) Giemsa stained blood smears were prepared for microscopic examination. Percentage (%) activity (parasite reduction) of each drug dose tested was determined using the following equation:

% activity = {(Parasitaemia in negative control - Parasitaemia in study group)/ Parasitemia in negative control} x 100.

#### Data Presentation and Statistical Analysis

The parasitaemia data from microscopy were analysed using R software. Boxplots were generated to portray the KO growth trends and parasite densities in presence and absence of the test drugs. Using the Student’s t-test in the Stata version 15.1 software, we computed means of percentage drug activity/percentage parasitaemia from each mouse in the treated group and compared them to that from mice in the control group.

#### Molecular docking

Homology modelling and structure validation. The sequences of *P. berghei* PM4, PM7, PM8, and Ddi1, referred to as receptors, were obtained from PlasmoDB (http://plasmodb.org/plasmo/) and their three dimensional (3D) structures predicted using SWISS-MODEL (28) available at https://swissmodel.expasy.org/. The best structures were downloaded, saved in PDB format and visualized using PyMOL (29). The quality of the modelled structures was validated using PROCHECK, for stereochemical evaluation (30), PRoSA-web, for detection of potential errors in the 3D structures (31) and verify_3D, for compatibility of the 3D structures with their respective amino acid sequences (32).

#### Ligand preparation

The structures of LP and SQ were downloaded from the ChemSpider available at http://www.chemspider.com/, an online database that provides access to unique chemical compounds (33). The CADD Group’s Chemoinformatics Tools and User Services, available at https://cactus.nci.nih.gov/translate/, was used to convert the chemical structures from Mol2 format into PDB format for compatibility with the docking software.

#### Docking with AutoDock Vina

Optimization of the grid box parameters of the receptors and the ligands was executed using ADT. The files were saved in the PDBQT format; their corresponding coordinates rewrote into a configuration file used for the docking strategy. Docking was carried out using Autodock Vina (34,35), for docking of a single ligand with a single receptor. The binding geometries (binding energies and the positional root-mean-square deviation; RMSD) of the ligands and the proteins were displayed on an output file. A ligand orientation with low binding energy indicates better affinity towards a receptor.

## Results

### Generation of the PM4, PM7, PM8 or Ddi1 knockout parasites: The Ddi1 gene was refractory to deletion thus essential for the growth of the asexual blood stage parasites

Overall, the transfection experiments yielded a total of three KO parasite lines. Of the three mice infected with schizonts transfected with PbGEM-039254 targeting the PM4, two mice had developed parasitaemia of >4% by day eight post parasite inoculation (7 days of pyrimethamine treatment). All the three mice infected with schizonts transfected with PbGEM-086320 and PbGEM-057101 targeting the PM7 and PM8 were parasite positive by day six post parasite inoculation. Three successive attempts to delete the Ddi1 failed to recover parasite twenty days post infection suggesting that the Ddi1 gene was refractory to deletion and thus essential for the asexual blood stage parasite growth. We then proceeded to genotype the PM4, PM7 and PM8 KO parasite lines using three sets of primers; QCR2/GW2, GT/GW1(2) and QCR1/QCR2. Both the WT and KO parasite genomic DNA (gDNA) was isolated and diagnostic polymerase chain reaction (PCR) amplification conducted. As expected, using the standard GW2 and the vector specific QCR2 primers, we obtained a fragment of 1kb, 0.8kb, and 1kb in PM4, PM7 and PM8 parasite lines (Fig 1) indicating the presence of the vector within the parasite. The PCR amplification using vector specific quality control primers, the QCR1 and QCR2 only amplified in the WT parasite line but failed to amplify in the KO lines confirming successful deletion of the respective genes. To further confirm correct integration of the specific vector into the correct chromosome and position, we utilized the standard primers GW1 or GW2 and the vector-specific primers GT. As expected, we obtained a PCR product of 2.9kb, 2.1kb and 2.6kb for PM4, PM7 and PM8 respectively. These results affirmed the successful generation of the KO parasite lines here referred to as PM4_KO, PM7_KO and PM8_KO. We used these parasite lines to assay drug response profiles with WT parasites as the reference line.

**Fig 1:**
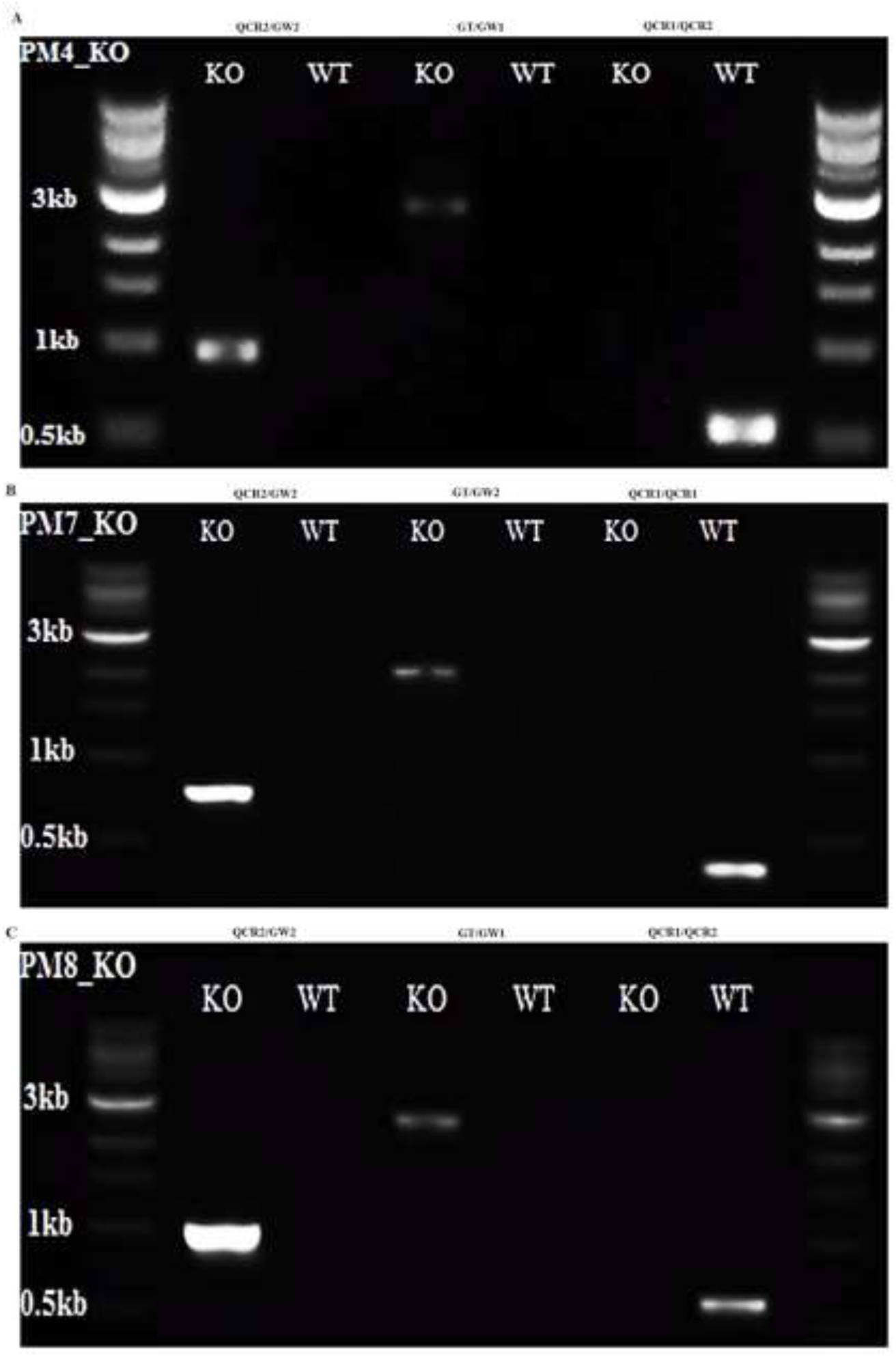
PCR genotyping of the knockout parasites; (A) PM4_KO (B) PM7_KO and (C) PM8_KO, and WT control lines.

The PCR amplification used three sets of primer pairs; QCR2/GW2, GT/GW1(2) and QCR1/QCR2. The PCR products were resolved in a 1% agarose gel electrophoresis. KO indicates the sizes of DNA fragments amplified from KO parasite lines while WT shows DNA fragment sizes from the control WT parasites. The amplification confirmed the presence of the vector, correct vector integration in the parasite genome and excision of the WT gene locus.

### The PM7 and PM8 genes are dispensable, deletion of the PM4 gene results in significant attenuation of growth rate of the asexual blood stage parasites

We evaluated the effect of the gene KO on the growth rate phenotypes by measuring the *in vivo* asexual stage parasite density (parasitaemia) for all the KO and WT parasites. The phenotype growth rate results revealed that deletion of PM4 significantly reduces normal parasite growth rate phenotype by 58% (*P* = 0. 0.003), suggesting a substantial contribution to the fitness of the asexual blood stage parasites. On the contrary, PM7_KO and PM8_KO exhibited significantly increased growth rate phenotypes compared to the WT parasites (*P* = 0.007 and *P* = 0.0007 respectively) D4 PI (Fig 2).

**Fig 2:**
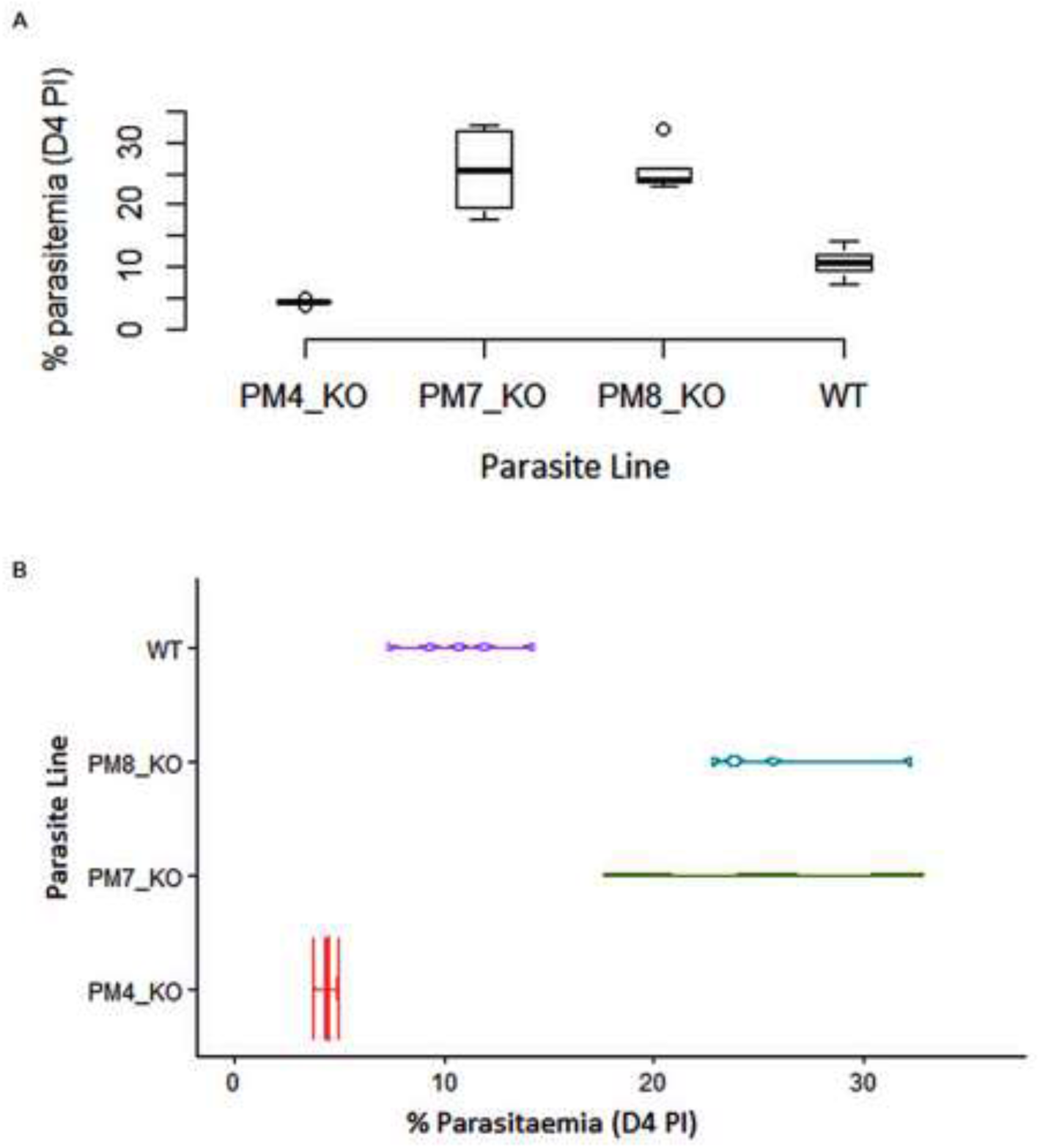
Deletion of the PM4 gene significantly attenuates the growth of asexual parasites. (A). Box plots showing the median percentage parasitaemia in mice infected with the PM4_KO, PM7_KO or PM8_KO parasite line relative to the wild-type (WT) parasite line as measured on day 4 post parasite inoculation in the standard 4-day suppressive test. (B). Violin plots showing the distribution of the parasites in mice infected with the PM4_KO, PM7_KO or PM8_KO parasite line relative to the wild-type (WT) parasite line as measured in the standard 4-day suppressive test. The PM4_KO parasites acquired a reduced growth rate phenotype (*P* = 0. 0.003), while the PM7_KO and PM8_KO parasites lines attained an increased growth rate phenotype.

### The PM7_KO and PM8 KO lines remained sensitive to LP and SQ while the PM4_KO line lost significant susceptibility to both LP and SQ

We assayed whether the deletion of the PM4, PM7 or PM8 affected susceptibility to LP, SQ, PQ and AQ. We reasoned that if the PM4, PM7 or PM8 are targets for the LP and SQ drugs, the KO parasite lines would lose susceptibility to the drugs. We determined the parasite susceptibility by calculating the activity of LP/RT at 40mg/kg.10mg/kg, SQ/RT at 50mg/kg.5mg/kg, AQ at (2.5 mg/kg and 1.25mg/kg) and PQ at (2.5mg/kg and 1.25mg/kg) on day four post parasite infection. The LP/RT significantly reduced the PM4_KO mutants’ parasitemia by 5.01% compared to a 15.71% in the WT parasites (*P* = 0.036). Similarly, the SQ/RT reduced PM4_KO mutants’ parasitemia by 5.29% compared to a 14.12% against the WT parasites (*P* = 0.030) (Fig 3B). However, the activity of both the LP/RT and SQ/RT against the PM7_KO and PM8_KO parasites were comparable to the percentage activity of WT parasites (Fig 3C and 3D). As expected, AQ and PQ were active against the WT parasites. At the 2.5mg/kg and 1.25mg/kg of AQ or PQ, the mutant parasite remained equally susceptible.

**Fig 3:**
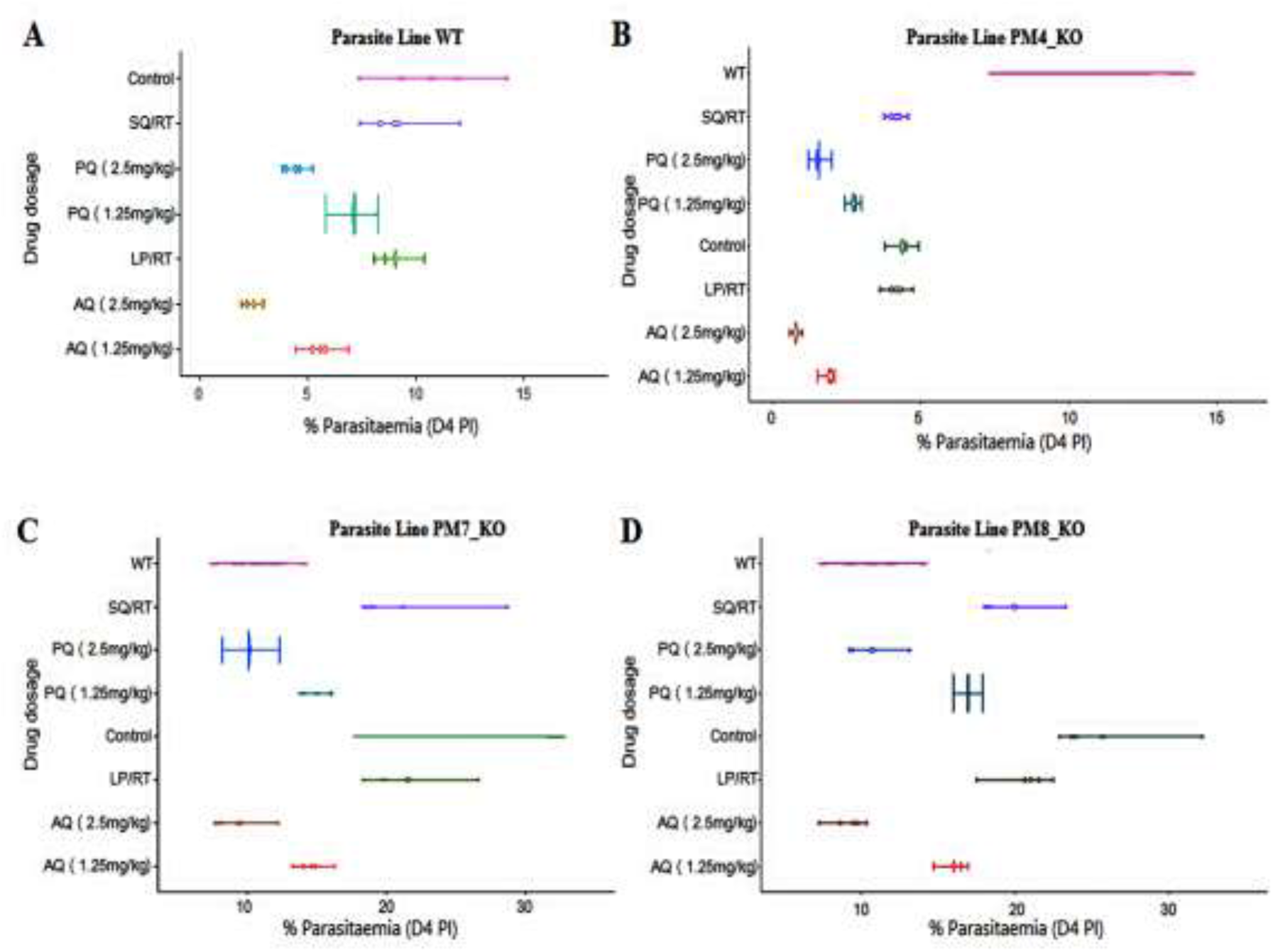
The PM4_KO parasites lost significant susceptibility to both LP and SQ. Violin plots showing growth profiles and the distribution of the asexual parasites. (A) The wild type (WT) parasite line displayed normal growth pattern and the expected susceptibility to amodiaquine (AQ), piperaquine (PQ), LP/RT and SQ/RT. (B) The PM4_KO parasite line exhibited a significant reduction in the growth outline and a significant loss of susceptibility to LP/RT (*P* = 0.036) and SQ/RT (*P* = 0.030) but not to AQ or PQ drugs (C) The PM7_KO and (D) PM8_KO parasite lines displayed rapid growth pattern as compared to the WT parasite line but retained susceptibility to AQ and PQ as well as to the RPIs; LP/RT and SQ/RT.

### Both the LP and SQ exhibited higher binding affinity for PM4 protein than PM7, PM8 or Ddi1

To estimate the degree of correctness for all the modeled structures, the Z-scores from PROSA-web server were determined and the proteins scored -5.09, -9.11, -7.94 and -7.52 for Ddi1, PM4, PM7 and PM8 respectively. The Z-scores implied that the structures of the modeled proteins were within the range of scores typically found with experimentally defined proteins. The averaged 3D-ID values were obtained from VERIFY 3D while PROCHECK confirmed residue positioning in the 3D structures and the values confirmed the integrity of the modeled structures (Table 2).

**Table 2:**
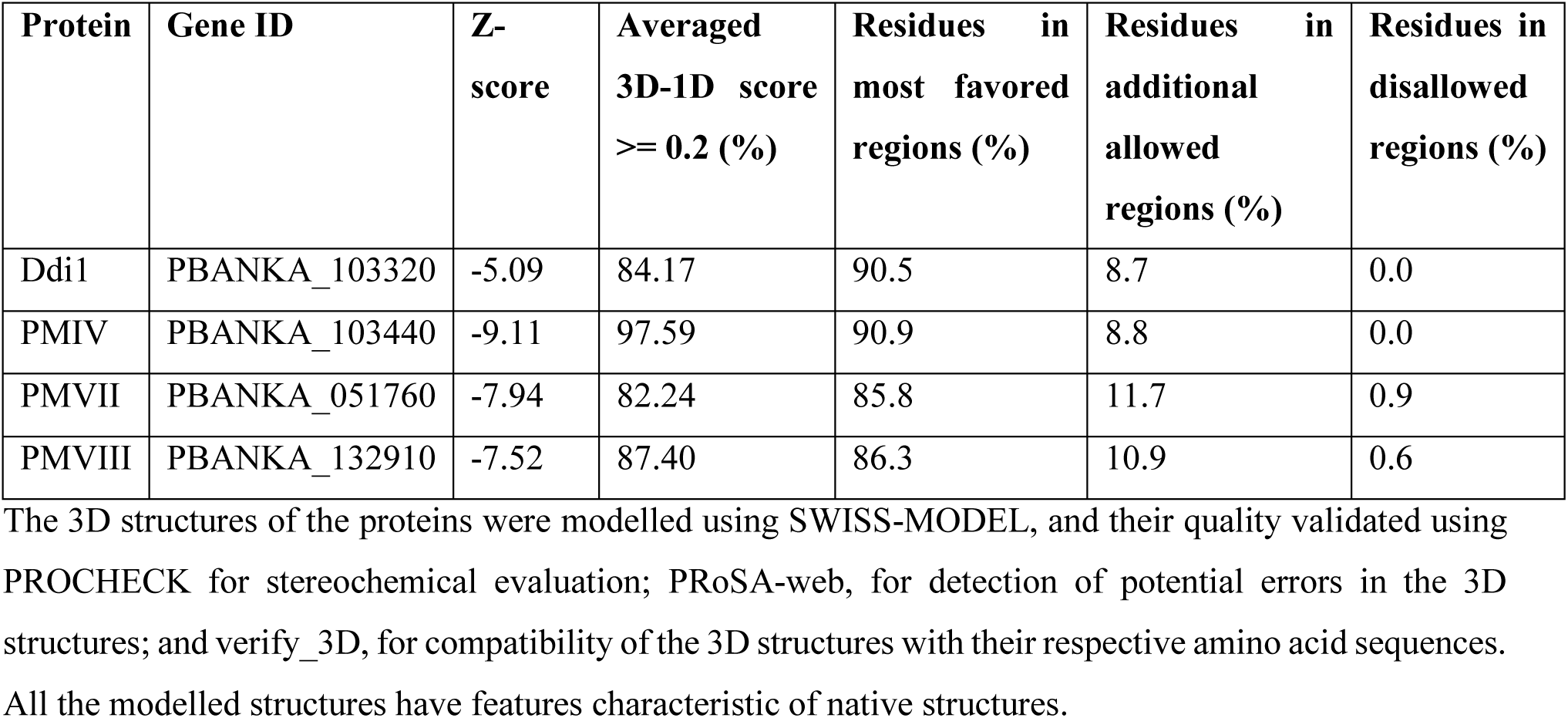
Validation of the modeled 3D structures of the *P. berghei* aspartyl proteases

The 3D structures of the proteins were modelled using SWISS-MODEL, and their quality validated using PROCHECK for stereochemical evaluation; PRoSA-web, for detection of potential errors in the 3D structures; and verify_3D, for compatibility of the 3D structures with their respective amino acid sequences. All the modelled structures have features characteristic of native structures.

When we evaluated the protein binding affinity, we determined the ligand conformations using the AutoDock Tools (ADT; Fig 4), and the best binding modes (mode 1). All the protein binding complexes showed high binding affinities, with the highest affinity at -10.6 kcal/mol and the lowest at -6.3 kcal/mol. Unlike the Ddi1 that had an equal binding affinity as HIV-1 asp protease for LP (-6.3 kcal/mol), the binding affinities of LP or SQ to PM4, PM7 or PM8 were higher than the affinity of the two drugs to the HIV-1 aspartyl protease (-6.3 kcal/mol). The PM4 had the lowest binding energy; -9.3 and -10.6 kcal/mol for LP or SQ respectively, indicating better binding with the LP and SQ (Figs 4 and 5).

**Fig 4:**
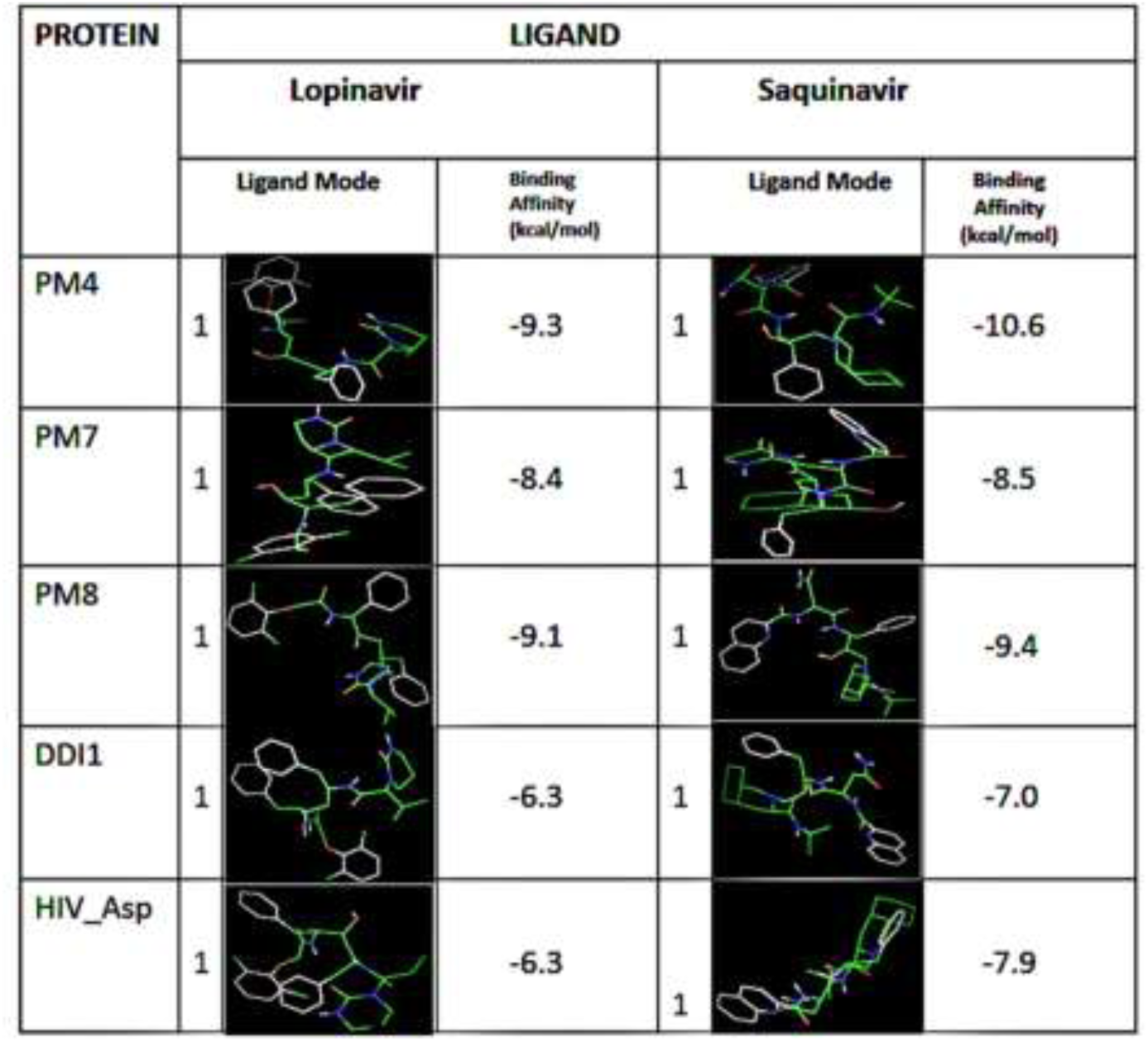
Both the LP and SQ exhibited high binding affinity for the PM4 protein. The binding geometries for docking of a single ligand with a single receptor as modelled using the Autodock Vina. The binding energies for LP and SQ to the PM4 were the lowest suggesting better affinity to the protein. The LP and SQ yielded higher binding affinity to both PM7 and PM8 as compared to the known RPIs target; the HIV Aspartic protease (HIV Asp). Both LP and SQ exhibited equal or low binding energy to the essential Ddi1 protein as compared to the HIV Asp.

**Fig 5:**
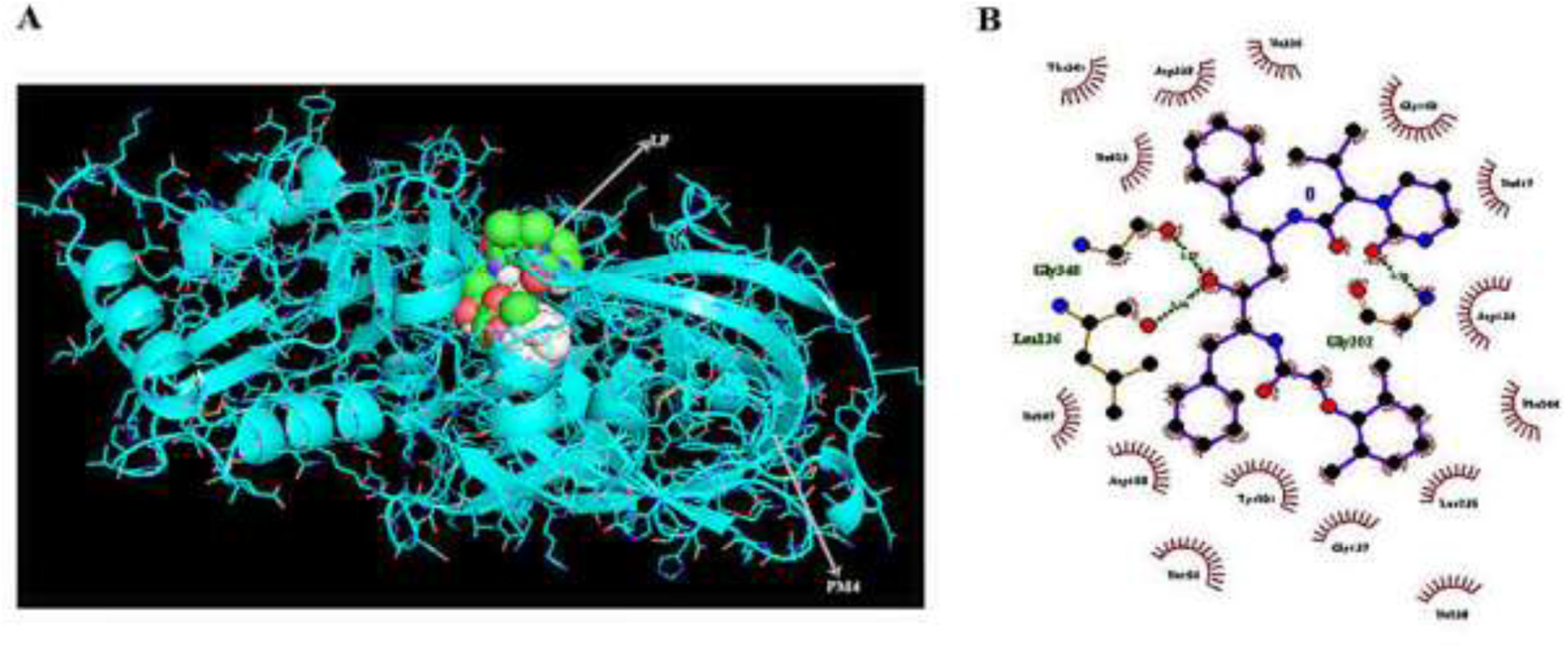
(A) The crystallographic structure of PM4 protein in complex with the lopinavir as visualized using PyMOL. (B) 2D LP-PM4 interaction diagram with the oxygen atoms shown in red, nitrogen atoms shown in blue while hydrogen bonds are shown as olive green dotted line. The interaction plots were generated using LigPlot+.

### The Ddi1 has a pepsin_retropepsin-like domain conserved across all *Plasmodium* species

We then reasoned that since deletion of the Ddi1 seemed toxic to the growth of the asexual blood stage parasites, the protein might have unique and conserved motifs across the different malaria parasites. We searched and analysed conserved motifs in the Ddi1 gene sequence using the Batch CD-Search (36) in the Conserved Domain Database (CDD). The CD-search results revealed that Ddi1 gene has a pepsin_retropepsin-like domain, with a catalytic motif, within its sequence. Interestingly, this motif is conserved in all the eight *Plasmodium* species that compared in our analysis (Fig 6).

**Fig 6:**
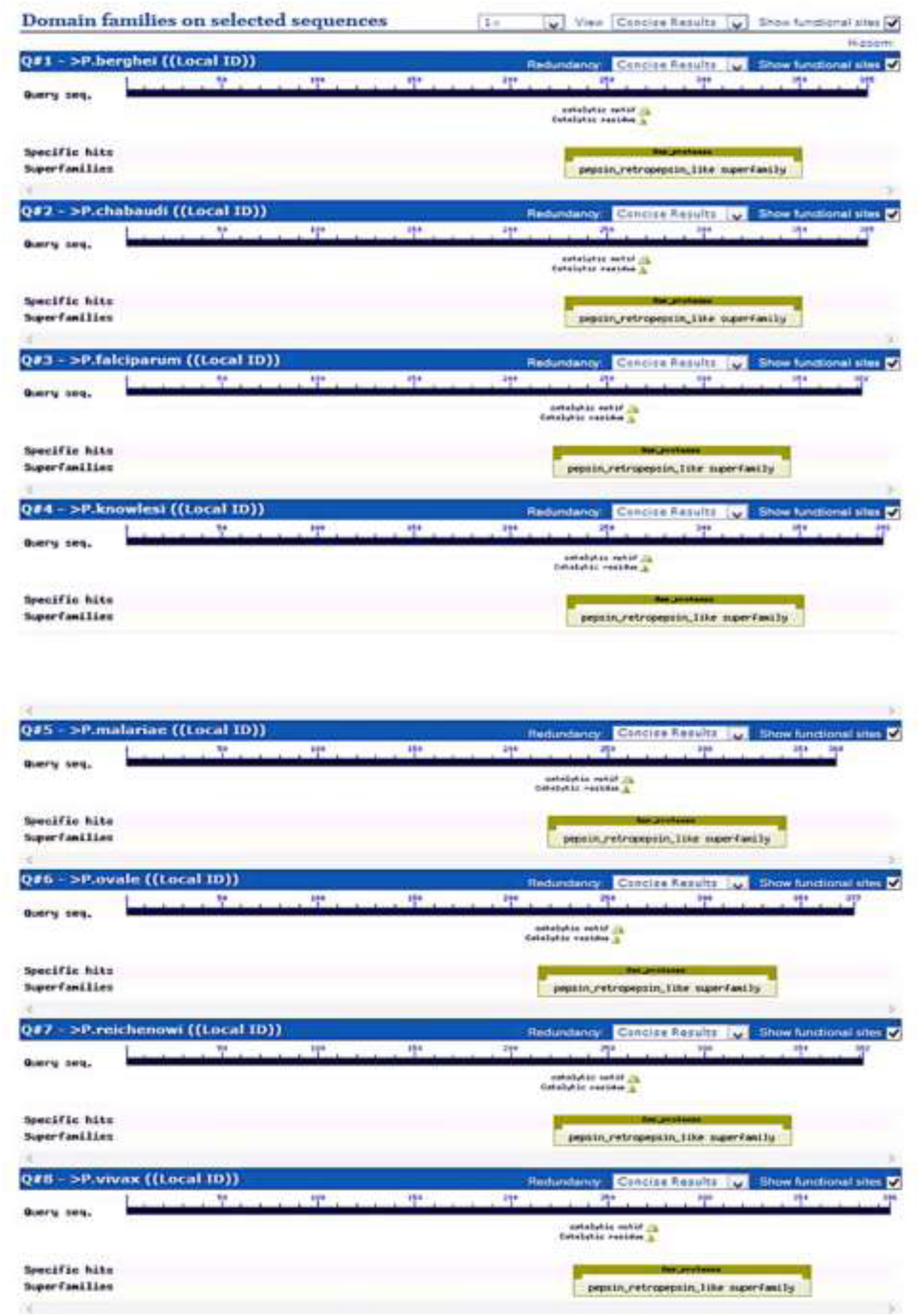
The Ddi1 protein has a conserved catalytic motif in its sequence. The Batch CD-Search showing the pepsin_retropepsin-like domain, with a catalytic motif conserved among the *Plasmodium* parasite species. The search involved eight *Plasmodium* species and all of them show the conserved motif between amino acid positions 250 and 280

## Discussion

The emergence and fast-spreading resistance against artemisinin-based combination drugs have spurred the identification and validation of novel antimalarial targets. Proteases are proven drug targets as evidenced by the use of protease inhibitors such as LP and SQ for the treatment of HIV/AIDS. This study considered both *Plasmodium* pepsin-like proteases; PM4, PM7 or PM8 as well as a retropepsin-like protease (Ddi1) as potential targets for RPIs. We investigated the essentiality of PM4, PM7, PM8 or Ddi1 by creating KO lines and used the lines to understand the possible mechanisms of action of LP or SQ. We generated KO *Plasmodium berghei* parasites deficient in the PM4, PM7 or PM8 genes but the deletion of the Ddi1 gene seemed toxic to the growth of the asexual blood stage parasites. The domain search within the Ddi1 protein confirmed the presence of a conserved catalytic motif among all the *Plasmodium* parasites species. The Ddi1 protein belongs to the A2 family of proteases, a retropepsin-like protease and is an active and functional aspartyl protease in *Leishmania major* parasites (37). The protease is involved in a novel, ubiquitin-dependent proteolytic, cell cycle control and reduction of extracellular protein secretion in *Saccharomyces cerevisiae* (38). Recently, studies have shown that an ortholog of yeast Ddi1 in *L. major* and human can complement protein secretion phenotype in Ddi1-deficient *S. cerevisiae* cells and that RPIs forestall the resultant complementation phenotype (39). The retropepsin-like family of proteases includes retroviral aspartyl proteases, retrotransposons and DNA-damage-inducible proteins in eukaryotic cells. From this study, the high binding affinities of the Ddi1 to the LP coupled with its essential role in the growth of the asexual blood parasite suggest that chemical compounds that inhibit the Ddi1 may form next generation of novel drugs.

To study the role of other aspartyl proteases in mediating drug action, we used the generated KO lines (PM4_KO, PM7_KO, and PM8_KO) to investigate the effect of gene deletion on the parasite growth rate phenotypes. Although a recent study on the functional profile of *Plasmodium* genome deleted the genes in *P. berghei*, the effect of the gene deletion on drug susceptibility was not measured (40). In agreement with this study, we have demonstrated that, whereas the PM7 and PM8 genes are dispensable during the asexual stages of parasite development, they exhibit an indirect interaction with asexual stage expressions. The loss of PM7 and PM8 genes increased the growth rate of asexual blood stage parasites suggesting that the deletion confer a growth advantage to the KO parasite. At D4 PI, the parasitemia for the PM7_KO and PM8_KO parasites was 2.5-fold higher than that of the WT parasites. On further scrutiny, we observed that, despite the PM4 being a dispensable gene, its deletion significantly reduces parasite asexual stage growth rates, Fig 2. The parasitemia for PM4_KO parasites was 2-fold lower than that of the WT parasites, suggesting a substantial contribution to the fitness of the parasite within the red blood cells. Whereas *P. falciparum* expresses four digestive vacuoles (DV) PMs; (PM 1, 2, histo-aspartic protease; HAP and PM4), there is only one identified DV PM in *P. berghei*, PM4. Therefore, successful deletion of PM4 in *P. berghei* suggests that the parasite could be relying on other enzyme pathways for haemoglobin degradation that sustains the supply of the amino acids for the intraerythrocytic parasite growth. The findings of PM4 deletion in *P. berghei* are consistent with the findings from (18), who documented that a quadruple KO of all the DV PMs (PM1, PM2, HAP, and PM4) in *P. falciparum* does not entirely impair parasite growth. However, the effect of the quadruple gene KO on drug susceptibility was not determined.

Malaria parasite haemoglobin degradation is a vital process during intra-erythrocytic parasite development. The plasmepsins, aminopeptidases, metalloprotease, falcilysin, and falcipains are found in the DV and are involved in degrading the haemoglobin. The presence of multiple enzyme pathways with overlapping specificity and function is strategic to increase parasite fitness (41,42). As earlier alluded, the deletion of PM4 confers significant growth disadvantage, and the KO parasite is less susceptible to LP and SQ. We thus argue that since the PM4 participates in the haemoglobin degradation (22,43), then the protein may be a target for both the LP and SQ. However, the presence of alternative enzymes for haemoglobin degradation provides the asexual blood stage parasites with an advantage albeit with cost in the growth rate.

Our drug profile results show that the activity of all the tested drugs against PM7_KO and PM8_KO parasites was nearly equivalent to the WT parasites. The activity of LP/RT and SQ/RT ranged between 15% and 21% against both the mutant parasites and WT parasites. These findings augment the fact that the asexual parasites do not express the PM7 and PM8 genes within the erythrocytes (42,43). Our study focused on the asexual blood stage parasites; both the LP and SQ are predicted to act on the parasite within the erythrocytes, thus although the LP and SQ may bind PM7 and PM8, the inhibition effect may not be observed in the asexual parasites within the erythrocytes. Indeed, studies involving PM8 have shown that it plays an essential role in the sporogonic stages of malaria parasite development (44). On the contrary, compared to the WT parasites, the LP/RT or SQ/RT activity against PM4_KO lines was 3-fold lower, demonstrating a potential role in the LP and SQ action, Fig 3. The PM4 is the only DV specific PM found in all *Plasmodium* species suggesting an essential role for the protein in the growth of the asexual blood stage parasite (45).

The modeling and molecular docking studies revealed that LP or SQ have high binding energies towards the *Plasmodium* aspartyl proteases. In fact, the drugs’ binding affinity towards PM4, PM7, PM8 or Ddi1 were higher than the affinity towards the HIV-1 aspartyl protease. However, because parasites contain many proteases with functional redundancy, unlike retroviruses, the *in vivo* activity of LP and SQ against the parasites is not proportional to the *in silico* binding affinities. The high binding affinity of LP or SQ towards PM4 correlates with the reduced sensitivity of PM4_KO parasites to the drugs. However, because of the redundant enzyme system involved in haemoglobin digestion for parasite asexual stage development, inhibition of PM4 does not entirely deprive of parasite survival. The high binding affinity shown by LP or SQ towards PM7 or PM8 (-8.4 kcal/mol to -9.4 kcal/mol) does not correlate with the *in vivo* drug suppression of asexual stage parasites probably because the enzymes are not essential and thus possibly not expressed during the erythrocytic asexual stages of parasite growth. Future work tailored towards understanding the functions of Ddi1 in *Plasmodium* parasites and its candidature as a target for drugs may inform the development of future drugs against the malaria parasite.

## Acknowledgments

We thank the team from the *Plasmo*GEM project at the Wellcome Trust Sanger Institute for providing the highly efficient gene modification resources used in this study. We also are grateful to Florence Ng’ong’a and Rosaline Macharia who assisted in the execution of the *in silico* work.

